# Gain control by sparse, ultra-slow glycinergic synapses

**DOI:** 10.1101/2021.05.03.442480

**Authors:** Varsha Jain, Laura Hanson, Santhosh Sethuramanujam, Ronald G. Gregg, Chi Zhang, Robert G. Smith, David Berson, Maureen A. McCall, Gautam B. Awatramani

**Affiliations:** Department of Biology, University of Victoria, Victoria, BC V8W 3N5 Canada; Departments of Ophthalmology & Visual Sciences, University of Louisville, Louisville KY 40202, USA; Anatomical Sciences & Neurobiology, University of Louisville, Louisville KY 40202, USA; Biochemistry & Molecular Genetics, University of Louisville, Louisville KY 40202, USA; Department of Neuroscience, University of Pennsylvania, Philadelphia, PA 19104, USA; Department of Neuroscience, Brown University, Providence, RI 02912, USA

**Keywords:** Glycine, glycine receptors, GABA, inhibition, direction selectivity, starburst amacrine cells, direction selective ganglion cell, retina

## Abstract

Retinal ON starburst amacrine cells (SACs) play a critical role in computing stimulus direction, partly in service of image stabilization by optokinetic nystagmus. ON SAC responses are sculpted by rich GABAergic innervation, mostly from neighbouring SACs. Surprisingly, however, we find that glycinergic narrow field amacrine cells (NACs) serve as their dominant source of inhibition during sustained activity. Although NAC inputs constitute only ∼5% of inhibitory synapses to ON SACs, their distinct input patterns enable them to drive glycine inhibition during the both light increments and decrements. NAC-to-ON-SAC inhibition appears to be mediated by ultra-slow non-canonical glycine receptors containing the α4 subunit, which effectively summate during repetitive stimulation. Glycinergic inhibition strongly decreases the output gain of the SACs, ensuring that their direction-selective output is maintained over their operating range. These results reveal an unexpected role for glycinergic pathways and receptor kinetics in modulating direction selectivity in the retina.

## Introduction

From as early as the work of Ramón y Cajal, it has been clear that inhibitory neurons form a more heterogeneous collection of types than do the excitatory neurons in many brain regions (Maccaferri and Lacaille, 2003). Advances in molecular, neurochemical and/or anatomical techniques have led to an explosion in the number of known interneuron types (Markram et al., 2004). Convergent evidence from such methods increasingly permits elaborate classification schemes, in which diverse inhibitory neurons are grouped into distinct, biologically meaningful types (Ascoli et al., 2008; DeFelipe et al., 2013). But to fully grasp the functional implications of interneuron diversity we need to develop a more complete understanding of their specific roles in intact and functioning neural circuits.

The retina is a particularly favorable region of the central nervous system for such an effort. More than 60 types of amacrine cells are present within single retinas, most with poorly understood functional roles (Yan et al., 2020). Here, we focus on inhibitory input to starburst amacrine cells (SACs), arguably the best known inhibitory interneurons of the retina with among the best-defined functional roles. SACs have occupied the spotlight mainly because of their unique ability to compute the direction of moving objects. Their radiating dendrites are the first elements in the visual pathway where direction selectivity is observed. Directional information computed in SAC dendrites is relayed through direction-selective ganglion cells (DSGCs) to sub-cortical/cortical visual areas, where it contributes to reflexive behaviors (e.g. image stabilization through optokinetic nystagmus; OKN), as well as perception (Rasmussen and Yonehara, 2020). Silencing SACs by pharmacological, genetic or pharmacogenetic means disrupts direction selectivity in downstream direction-selective ganglion cells (DSGCs) and OKNs, underscoring the critical role for SACs in this computation (Taylor and Smith, 2012) (Wei, 2018).

Recent studies have made great strides in understanding the precise anatomical and functional properties of connections to SACs, providing valuable insights into how SACs compute direction. In terms of the inhibitory inputs to ON SACs, which we focus on in this study, serial block-face electron microscopic (SBEM) reconstructions show that the vast majority of inhibitory contacts are GABAergic, arising mainly from neighboring SAC and a few from wide-field amacrine cells (WACs) (Briggman et al., 2011; Ding et al., 2016). Mutual GABAergic inhibition between neighboring SACs has been proposed to contribute to direction selectivity through a bias toward connectivity between anti-parallel SAC dendrites (Chen et al., 2016; Ding et al., 2016; Lee and Zhou, 2006; Munch and Werblin, 2006). On the other hand, WACs have been shown to mediate a TTX-sensitive surround suppression, which contributes to the specific size selectivity and the contextual modulation of the neuronal responses within the circuit (Hoggarth et al., 2015; Huang et al., 2019).

Ding et al. also found that a small percentage of inhibitory input to ON SACs (<5%), comes from a glycinergic narrow-field amacrine cells (NACs) (Heinze et al., 2007). Consistent with the paucity of NAC synapses (<5%), spontaneous and light-evoked inhibitory input to SACs are little affected by glycine receptor blockade but strongly affected by GABA antagonists (Lee and Zhou, 2006; Majumdar et al., 2009). Furthermore, glycine inhibition does not appear to affect direction selectivity in SAC dendrites (Hausselt et al., 2007), or in downstream DS ganglion cells (DSGCs), whose computation relies on cholinergic/GABAergic SAC output (Caldwell et al., 1978; Wyatt and Day, 1976). Together, these results suggest a minor role for glycinergic inhibition in the DS circuit, which is thus usually ignored (Wei, 2018).

However, local application of glycine evokes large currents in ON but not OFF SACs (Ishii and Kaneda, 2014; Majumdar et al., 2009). Moreover, GlyR blockade greatly increased light-evoked acetylcholine (ACh) release in classic bulk-release studies (Cunningham and Neal, 1983). To comprehensively address the role of glycinergic inhibition in the SAC circuit, here we characterized the anatomical properties of NAC circuitry in an existing SBEM dataset (Ding et al., 2016). In addition, we assessed the functional properties of the inhibitory pathways controlling SACs using pharmacology and a variety of mouse genetic knockout (KO) models (including a novel *Glra4* KO), in combination with electrophysiology, 2-photon ACh sensor and Ca^2+^ imaging techniques. We show that although glycinergic NACs comprise only 5% of all inhibitory input (**Figure 1A-B**) (Ding et al., 2016), they can dominate inhibition to SACs during repetitive stimulation, changing the way moving stimuli are processed according to the recent history of activity.

**Figure 1:**
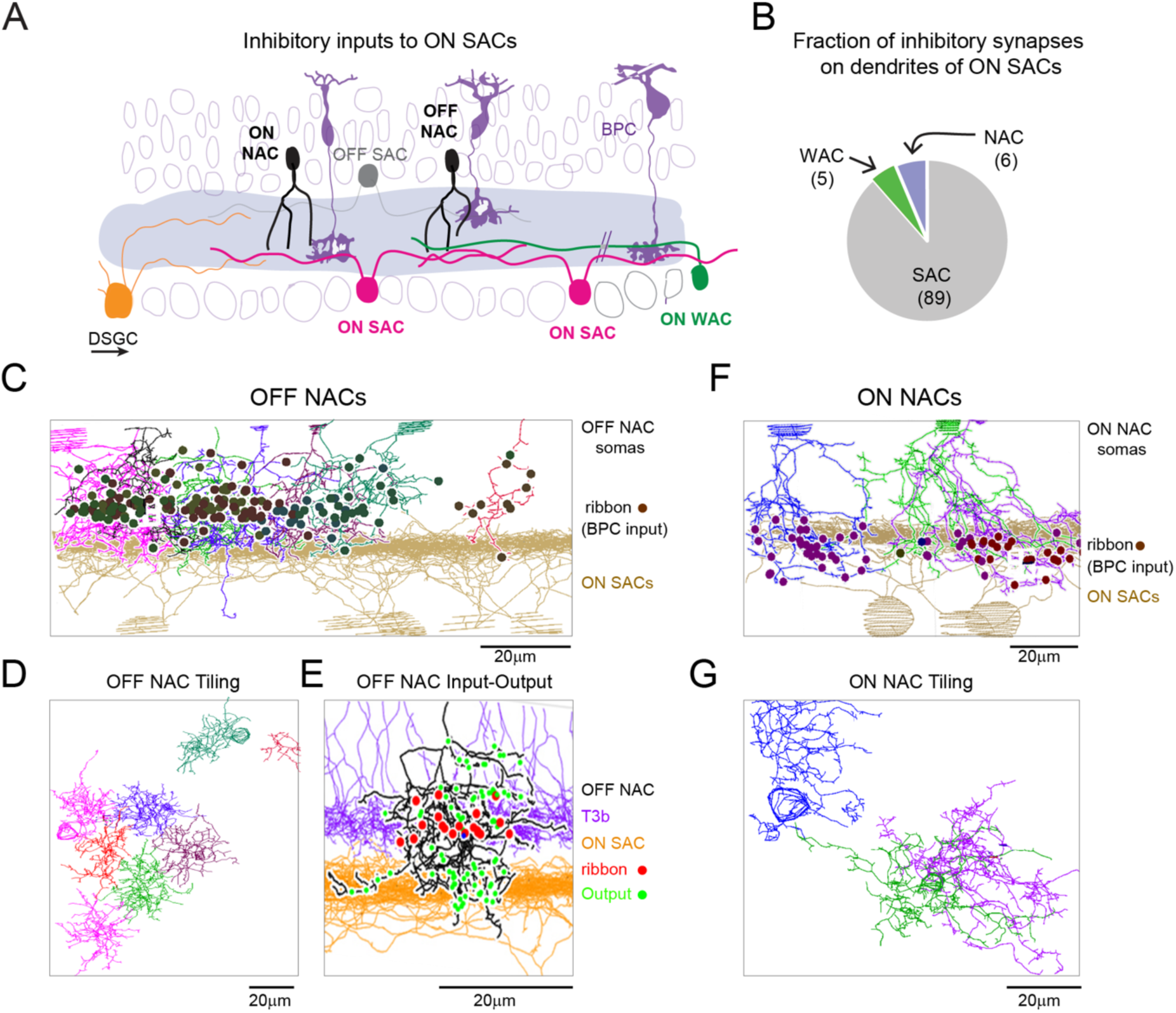
ON and OFF type of narrow field amacrine cells provide inputs to ON SAC dendrites. **A.** Three types of inhibitory amacrine cells contact dendrites of the ON starburst amacrine cell (SAC), including neighboring ON SACs, wide- and narrow-field amacrine cells (WACs and NACs, respectively). SACs provide direction-selective (DS) inhibition to neighboring SACs and to downstream DS ganglion cells (DSGCs). Bipolar cells (BPCs), driven by photoreceptors (not shown) are the source of excitation to SACs. **B.** NAC synapses constitute a small fraction of inhibitory synapses in ON SACs (adapted from (Ding et al., 2016)). Numbers in parentheses denote the percent of synapses from these cell types onto ON-SACs. **C-E**. Reconstructions of OFF NACs (n = 8; colored cells) identified by back tracing non-ribbon containing synapses on SAC dendrites (brown) in an SBEM dataset in which ultrastructure is preserved (Ding et al., 2016). Note, BPC input (ribbons; dots) are not in the ON SAC band. **D.** The axonal arbours of OFF NACs shown in C appear to form a non-overlapping mosaic, indicating the identified OFF NACs are of a single type. **E.** Back tracing ribbon-synapses (red dots) contacting OFF NAC (black) identifies mainly type 3b bipolar cells (purple). NAC output synapses (green) are more broadly distributed with a strong concentration in the ON SAC band (orange). **F-G.** A similar analysis also identifies ON NACs (colored cells; n=3) as presynaptic partners to ON SAC (brown). ON-NACs receive ribbon synapses (dots) from type 7 bipolar cells. **G.** The axon terminals of ON-NACs shown in F.

## Results

### ON and OFF NACs in the direction-selective circuit

We extended the earlier serial EM analysis of (Ding et al., 2016), using the same dataset, to provide a more complete account of the narrow-field amacrine-cell types (NACs) thought to provide glycinergic input to ON SACs. (**Figure 1**). In agreement with the earlier study (**Figure 1B**), were found NACs provided only a small minority of all inhibitory inputs. These NACs could be grouped into two distinct types based on the stratification of their processes, their patterns of bipolar synaptic input, and the independent, tiling mosaics formed by their processes (**Figure 1C, E, F**). The first is an OFF NAC, apparently equivalent to AC Type 21 of (Helmstaedter et al., 2013) (**Figure 1C-E**). Though their processes extend profusely into both ON and OFF sublayers of the IPL, their excitatory ribbon synaptic input appears to come almost exclusively from Type 3b OFF cone bipolar cells in the OFF sublayer (30 ribbons; 27 T3b; 2 T4; 1 T2). Their outputs, however, are distributed in both the ON and OFF sublayers, and there is a band of particularly dense output at the level of the ON SAC plexus (**Figure 1C, E**). (Figure 1E). Their dendritic fields are 30 ± 5 µm in diameter (N = 8) and form a clear mosaic.

The second type is an ON NAC, apparently corresponding to AC Type 22 (Helmstaedter et al., 2013). The ON NACs had small dendritic fields 40 ± 5µm (N= 3) but too few were reconstructed to determine if they form a clear mosaic (**Figure 1F, G**). Ribbon containing boutons (bipolar cell) synapsing with ON NACs were almost exclusively in the ON IPL sublayer, arising overwhelmingly from T7 and T6 cone bipolar cells (**Figure 1F**), especially T7 (among identified ribbon inputs to 3 ON NACs : T7 (n=43); T6 (n=23); with no more than three ribbons each found from T3b (n=2), T5i (n=3), T8 (n=1) and T9 (n=2)). The ON NACs also make synaptic contacts to diverse amacrine cells and retinal ganglion cells, but this connectivity was not characterized further. The input/output distributions of the ON and OFF NACs predict that they will drive weak glycinergic inhibition to ON SACs at the onset and offset of the light stimulus, respectively.

### ON SACs receive powerful glycinergic inputs

Indeed, ON and OFF glycinergic IPSCs could be measured in ON SACs (V_HOLD_ ∼ 0mV, the excitatory reversal potential**; Figure 2A**). However, contrary to the anatomical representation, the glycinergic inputs were surprisingly large relative to the GABAergic input. In these experiments, SACs were targeted for voltage-clamp recordings in the Chat^cre^::Ai9 transgenic mouse retina using two-photon imaging, and stimulated with stationary spots (100 µm, 100% positive Weber contrast), which are not likely to stimulate WAC inputs. To minimize the GABAergic output of the recorded SAC onto neighboring SACs that could potentially result from holding the membrane potential at 0 mV, we included 1 mM EGTA in the recording pipette.

**Figure 2:**
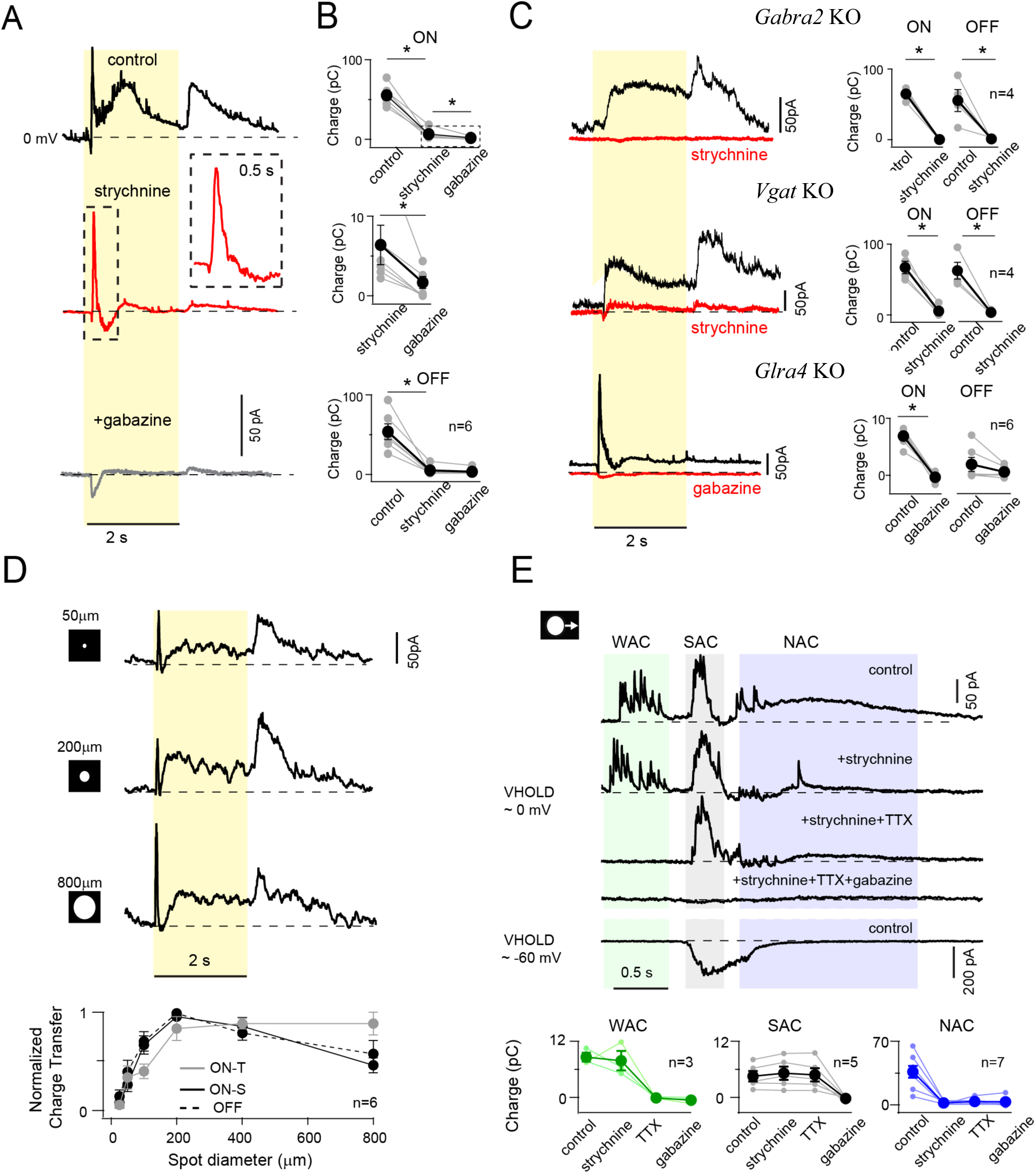
Glycine, not GABA receptors, mediate the dominant form of inhibition to SACs. **A.** IPSCs recorded from an ON SAC (at ∼0mV) evoked by small stationary spots (2 s;100 μm diameter) in control (black), in the presence of GlyR antagonist (1 µM strychnine, red) and following addition of the GABA_A_R antagonist, gabazine (grey). Inset shows the transient response in the boxed region at higher magnification. **B.** Average ON (*top and middle*) and OFF (*bottom*) inhibitory responses measured across 6 cells (black-mean ± SEM, grey-individual cells) in three conditions as shown in A. The middle panel plots the same data as the top panel and is magnified to highlight the significant reduction of ON inhibitory charge in gabazine. (* p < 0.05, t-test). **C.** ON SAC IPSCs measured in Gabra2 KO mouse (GABA_A_ α2 receptors are knocked out in SACs) and Vgat KO mouse (Vesicular GABA transporter is knocked out from SACs) lack transient ON component. IPSCs in these mice are completely blocked by strychnine (*left-top and middle*). By contrast, SAC IPSCs in *Glra4* KO mice mouse exhibited an ON transient responses (but not sustained responses), which are blocked by gabazine. *Right* ON and OFF responses measured for a population of cells in different KO mice (mean ± SEM, *p<0.001). **D.** ON SAC IPSCs evoked by stationary spots of increasing diameter. *Bottom-* Normalized ON transient (ON-T), ON sustained (ON-S) and OFF responses are plotted as a function of spot size. **E.** IPSC evoked by a moving spot typically have multiple components, each dominated by a single source (WAC, SAC and NAC) with sensitivity to different drugs (as indicated). EPSC (V_hold_ ∼ −60 mV, last trace) measured in control condition provides an indication of the excitatory receptive field for reference. The average responses (mean ± SEM) for three inhibitory components in different drugs quantified across a population of ON SACs.

Under these conditions, the SAC IPSC evoked by the onset of the stimulus comprised of a brief transient followed by a sustained phase, while the response to the light offset was purely sustained (**Figure 2A, B**). Bath application of the GlyR antagonist (1 µM strychnine) significantly suppressed the sustained component of the ON IPSCs and completely blocked the OFF IPSCs. The residual transient ON component was abolished upon additional application of the GABA_A_ receptor antagonist (5 µM SR-95331: gabazine). These results are consistent with the idea that the transient ON input is mediated by GABAergic SACs, whereas the sustained ON/OFF IPSCs are mediated by glycinergic NACs. For simplicity we will refer to the transient ON GABAergic IPSCs as the SAC component, and the sustained ON/ OFF IPSCs as the NAC component.

The origins of transient and sustained forms of inhibition were also assessed using a variety of KO mouse models. Specifically, we examined SAC IPSCs in two mouse lines in which mutual SAC-SAC GABAergic inhibition is selectively suppressed in distinct ways. In the first mutant line, SAC-SAC GABAergic inhibition was compromised postsynaptically, by conditionally deleting the α2 subunits of GABA_A_ receptors in SACs (Gabra2^fl/fl^ x ChAT^cre^) (Chen et al., 2016). In the other line, SAC GABA release was reduced by the deletion of the GABA vesicular transporter (Vgat^fl/fl^ x ChAT^cre^) (Bleckert et al., 2018; Pei et al., 2015). In both these KO mice lines, the light-evoked inhibitory responses measured in SACs lacked the transient ON IPSC component, confirming that they were mediated by SACs (**Figure 2C**). Additionally, the sustained ON and OFF IPSCs that were observed in these KOs, were eliminated by application of strychnine (**Figure 2C**). On average, the strength of the NAC ON and OFF IPSCs measured in these KO SACs were comparable to that measured in wild-type SACs (**Figure 2C**). These results confirm that GlyRs mediate a slow ON/OFF postsynaptic inhibition to ON SACs, while GABA_A_Rs mediate a fast, transient ON inhibition.

Previously, based on the prominent colocalization of GlyR*α*4 receptors with ON SAC cholinergic bands, along with a functional analysis of miniature IPSCs in a variety of *Glra* KO SAC (GlyRa1, a2 & a3 KOs) it has been suggested that glycinergic inhibition in ON SACs is in part mediated by GlyR*α*4-containing subunits (Heinze et al., 2007; Majumdar et al., 2009). To directly test this hypothesis, we generated a *Glra4* KO mouse and validated the absence of GlyRα4 expression using immunohistochemistry (**Supplementary Figures 1 and 2**). Indeed, *Glra4* KO SACs lacked sustained ON and OFF inhibitory currents. The transient SAC components in these mutant SACs was similar in strength and kinetics to SAC mediated inhibition measured in wild-type retinas in the presence of strychnine (6 ± 2 pC; p > 0.1) **(Figure 2C)**. Gabazine completely blocked the transient ON IPSCs in *Glra4* KO SACs, confirming that GABAergic SAC-SAC inhibition remains intact in the absence of GlyR*α*4 expression. Taken together, these results support the proposal made more than a decade ago: *α*4-containing GlyR subunits mediate glycinergic inhibition in SACs (Heinze et al., 2007; Majumdar et al., 2009).

Next, we tested the spatial extent over which glycine inhibition is recruited, by evoking IPSCs in ON SACs using spots of different diameters (**Figure 2D**). Both the ON and OFF sustained glycinergic IPSCs had a clear centre-surround organization (**Figure 2D**). The centre was ∼ 200 µm in diameter, which matches the spatial extent over which NACs are anatomically connected to SAC dendrites (Ding et al., 2016). However, the ON transient GABAergic IPSCs did not vary with spot size. This is likely to be due to a combination of decreases in SAC-SAC inhibition and increases in WAC inhibition as a function of increasing spot size (Hoggarth et al., 2015).

Interestingly, in response to moving stimuli (200 µm diameter spot, moving at 1000 µm/s), IPSCs appeared to have multiple temporally dispersed components, reflecting the contributions of WACs, SACs, and NACs (**Figure 2E**). The first component was evoked when the stimulus was 500-700 μm away from the recorded SACs dendritic field and could be blocked by either gabazine or 1µM TTX, consistent with the idea that it is mediated by WACs (note, TTX does not eliminate the output of non-spiking SACs or NACs (Hoggarth et al., 2015). However, the WAC component was not consistently observed with small spot stimuli, and we did not analyze its contributions further in this study. The next IPSC component was brief, gabazine-sensitive, TTX-insensitive, and was initiated contemporaneously with the EPSC. This component represents lateral GABAergic SAC inhibition, that helps shunt the initial excitatory inputs (**Figure 2E** bottom trace). Note, the initial excitatory inputs occur on the “null” SAC dendrite pointing in the opposite direction at the moving stimulus (see Ding et al., 2016). The third IPSC component was sustained and activated only after the stimulus left the SAC’s excitatory receptive field. This component was insensitive to gabazine and TTX, but strychnine-sensitive suggesting that it arises from NACs (**Figure 2E**). For moving stimuli, ON and OFF components were not apparent, likely because NAC IPSCs are extremely slow to activate. Thus, in the context of simple spot stimuli, SAC but not NAC inhibition appears to be important for controlling starburst function (Ding et al., 2016).

### Glycinergic unitary events exhibit slow kinetics

To understand the origin of the slow glycinergic IPSCs, we next examined the kinetics of synaptic glycinergic events that arise spontaneously in wild-type ON SACs. Under control conditions, spontaneous IPSCs (sIPSCs) varied widely in their kinetics and peak amplitudes (**Figure 3A**). Glycinergic sIPSCs, isolated in the presence of gabazine, were slow to rise and extremely slow to decay (τ_rise_ =2.8±1.2 ms; τ_decay_ = 64 ± 34 ms; n = 146 events from 5 cells; **Figure 3B**), perhaps amongst the slowest sIPSCs observed in the brain to date (also see (Majumdar et al., 2009)). Slow glycinergic sIPSCs were relatively small in amplitude (12 ± 4 pA), difficult to measure, and may contribute to a tonic current (see **Figure 4C**). By contrast, GABAergic sIPSCs, isolated in the presence of strychnine, were significantly larger (p<0.01) and faster (p<0.01) (**Figure 3B & C**). The decay kinetics of sIPSCs measured in strychnine were similar to those measured in *Glra4* KO ON SACs (p >0.05) (**Figure 3B & D**). Gabazine completely blocked the fast sIPSCs in 5/7 ON SACs. In the remaining two SACs, the residual sIPSCs measured in gabazine had intermediate kinetics, suggesting additional types of GlyRs may contribute to glycinergic synaptic inputs. These results indicate that transmission at unitary glycinergic synapses is extremely slow, likely arising from the biophysical properties of GlyR*α*4 subunits. While sIPSCs mediated by GlyRs are slow compared to those mediated by GABARs, they are significantly faster than the light-evoked glycinergic IPSCs. Thus, it is likely that vesicle release from NACs is slow and sustained. Together these factors results in a sustained inhibition that builds up and relaxes over many hundreds of milliseconds, outlasting the presence of the light stimulus.

**Figure 3:**
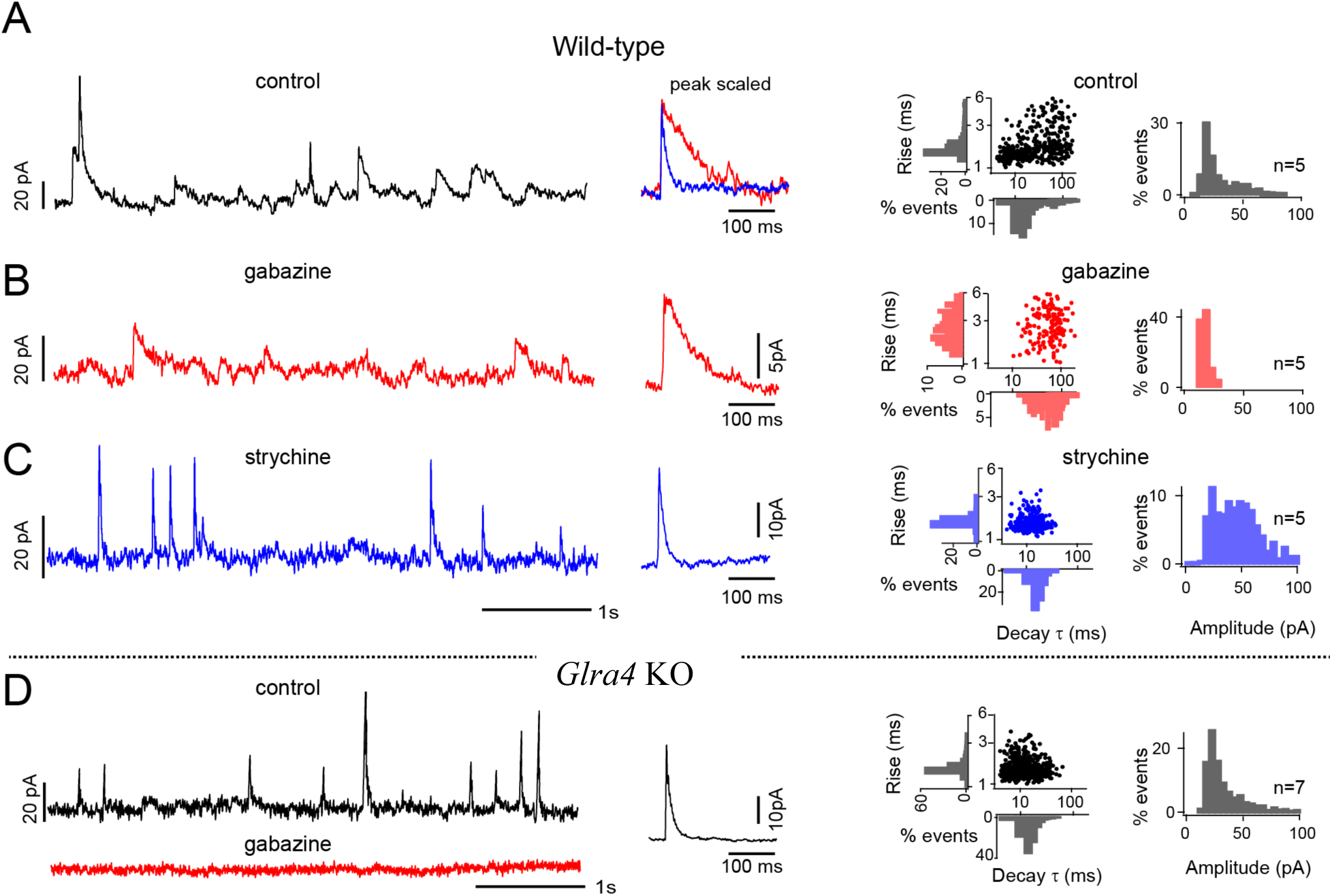
GlyR*α*4 receptors in SACs mediate slow and delayed sIPSCs. A. sIPSCs measured from ON SACs in wild-type mice at 0 mV in control (*left panel*). Normalized average fast (blue) and slow (red) sIPSCs (*middle panel*). Frequency distribution of rise, decay time and amplitude of the population events (*right panels*; n = 5 SACs). **B-C**. Same as A, but in gabazine (B) and strychnine (C) in wild-type SACs. **D.** same as A & B except SAC recordings were made in *Glra4* KO retina.

**Figure 4:**
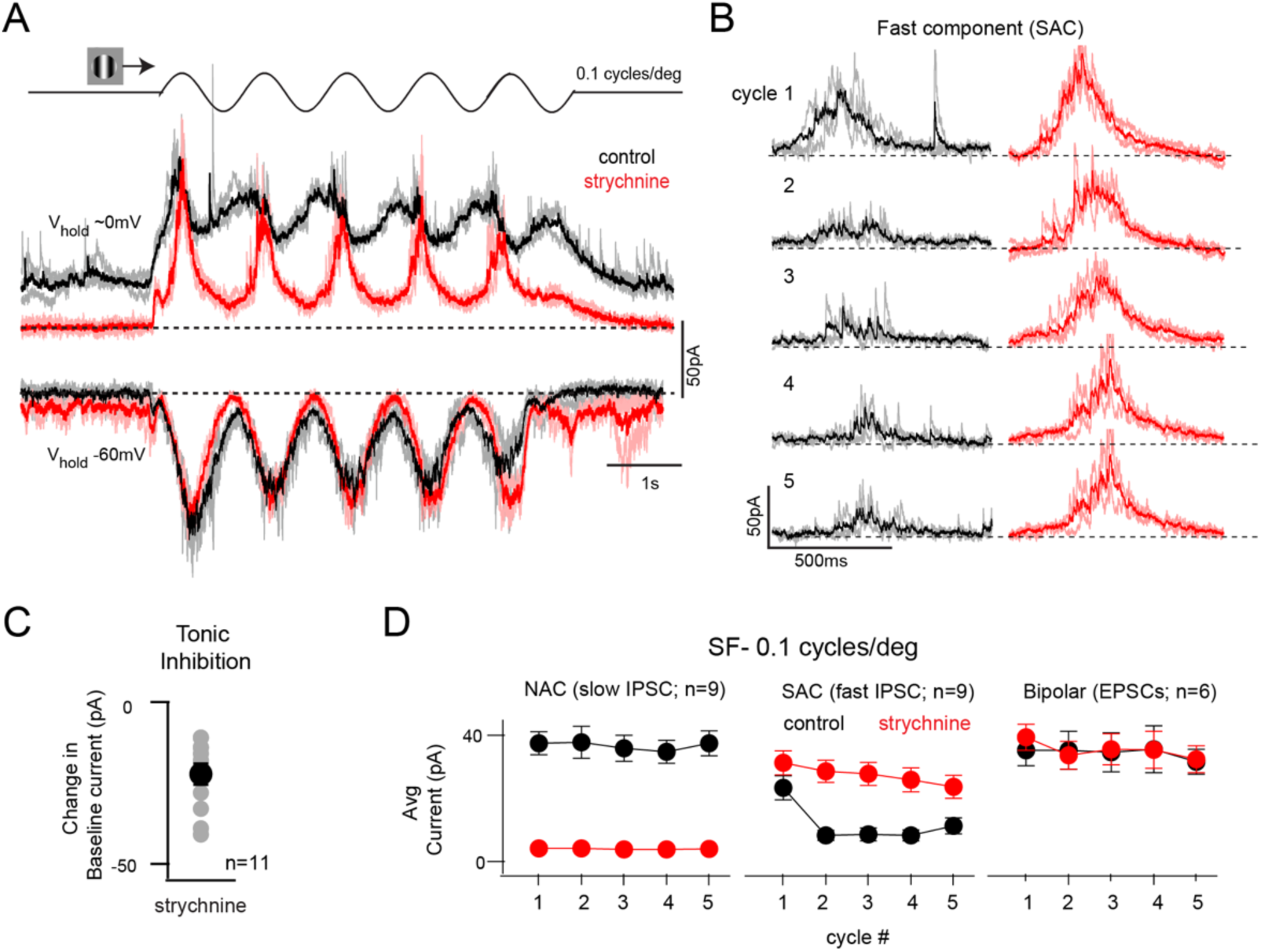
Glycinergic inhibition suppresses SAC output under sustained visual simulation. **A.** ON SAC IPSCs and EPSCs under control (black) and strychnine (red) evoked by drifting sinusoidal gratings (SF 0.1cycles/degree; TF 1Hz, indicated on top; also see Supplementary Figure 3 for responses evoked by 0.01cycles/degree) presented through a 500 μm diameter aperture. Dark traces are the average of 3 trials shown in a lighter shade. **B.** Transient IPSC evoked at each cycle of the stimulus, in control (black) and strychnine (red). Note, the slow NAC component has been subtracted for clarity. **C.** Net change in the baseline holding current after the application of strychnine. **D.** Strychnine has distinct effects on the NAC-(slow IPSC; left), SAC-mediated IPSCs (middle) and EPSCs (right) at each of the 5 cycles of the stimulus (control, black; strychnine, red). Data are represented as mean ± SEM.

### NACs control SAC output during sustained stimulation

In contrast to moving spot stimuli, when the retina was stimulated using drifting gratings we found NAC inhibition played an important role in controlling SAC output. In these experiments, gratings were presented through a 500µm diameter circular aperture to minimize WAC inhibition. Figure 4A illustrates responses to a grating which had a spatial frequency (SF) of 0.1 cycles/degree and temporal frequency (TF) of 1 Hz, showing that they were dominated by slow IPSCs (**Figure 4A & D**). The transient IPSCs were significantly smaller in amplitude and rode on top of the sustained inhibition. When the sustained phase was subtracted, it became evident that the transient IPSCs were strongly suppressed during the course of the stimulus (**Figure 4B;** see methods). The current averaged over cycles 2-5 was only ∼25-30% of the current evoked during the first cycle (**Figure 4B, D**).

Bath application of strychnine reduced a small tonic outward current (**Figure 4C**) as well as the sustained phase of the light-evoked IPSCs. It also greatly enhanced the transient SAC IPSC amplitude, indicating that under control conditions, NAC inhibition strongly supresses SAC output (**Figure 4A, D**). It is interesting to note that the initial response to the grating, appeared to be less affected by NACs. This is consistent with the finding that glycinergic inhibition is slow to activate (**Figure 4B, D**). Under GlyR blockade, the strength of the SAC-mediated IPSCs transient evoked at each cycle of the grating, did not change over the duration of the stimulus. And the phase advance of the response that occurred at each cycle was also less noticeable (**Figure 4B**). Similar effects of strychnine were observed for lower SF gratings (**Supplementary Figure 3**). In contrast to the IPSCs, strychnine did not significantly alter the SAC EPSCs measured at V_HOLD_ ∼ −60 mV, indicating that glycinergic modulation occurs mostly at the dendrites of ON SACs (**Figure 4A, D**).

One caveat in voltage-clamp experiments, is that the quality of the voltage-clamp is expected to change to some extent after blocking sustained inhibition, which would lead to underestimates of the relative contribution of the SAC component and exaggerate the degree to which SAC output is controlled by NACs. To independently verify the conclusions of our electrophysiological measurements we monitored light-evoked SAC output optically, using a G-protein-coupled receptor activation-based sensor (GRAB ACh 3.0 or simply ACh 3.0; kindly provided by Dr. Yulong Li, Peking University). Injecting AAV vectors to express Cre-dependent sensors intravitreally in Oxtr-Cre (**Figure 5A**) or Cart:Cre mice led to robust expression of the sensor 10-15 days after the injection (See methods) (Sethuramanujam et al., 2021). Stimulating the retina with drifting gratings produced reliable changes in ACh3.0 fluorescence in SAC or DSGC dendrites (**Figure 5B, C**). Brief exposure to strychnine doubled the ACh3.0 response (**Figure 5B-D**), and these effects were reversed when the drug was washed off. However, unlike our electrophysiological results in many cases the initial ACh sensory response was also augmented. This is likely because NACs activated by the laser-scanning prior to the presentation of the gratings, produced tonic glycinergic currents. Together with results from our electrophysiological experiments, these data demonstrate that glycinergic inhibition suppresses the SAC output during sustained visual stimulation.

**Figure 5:**
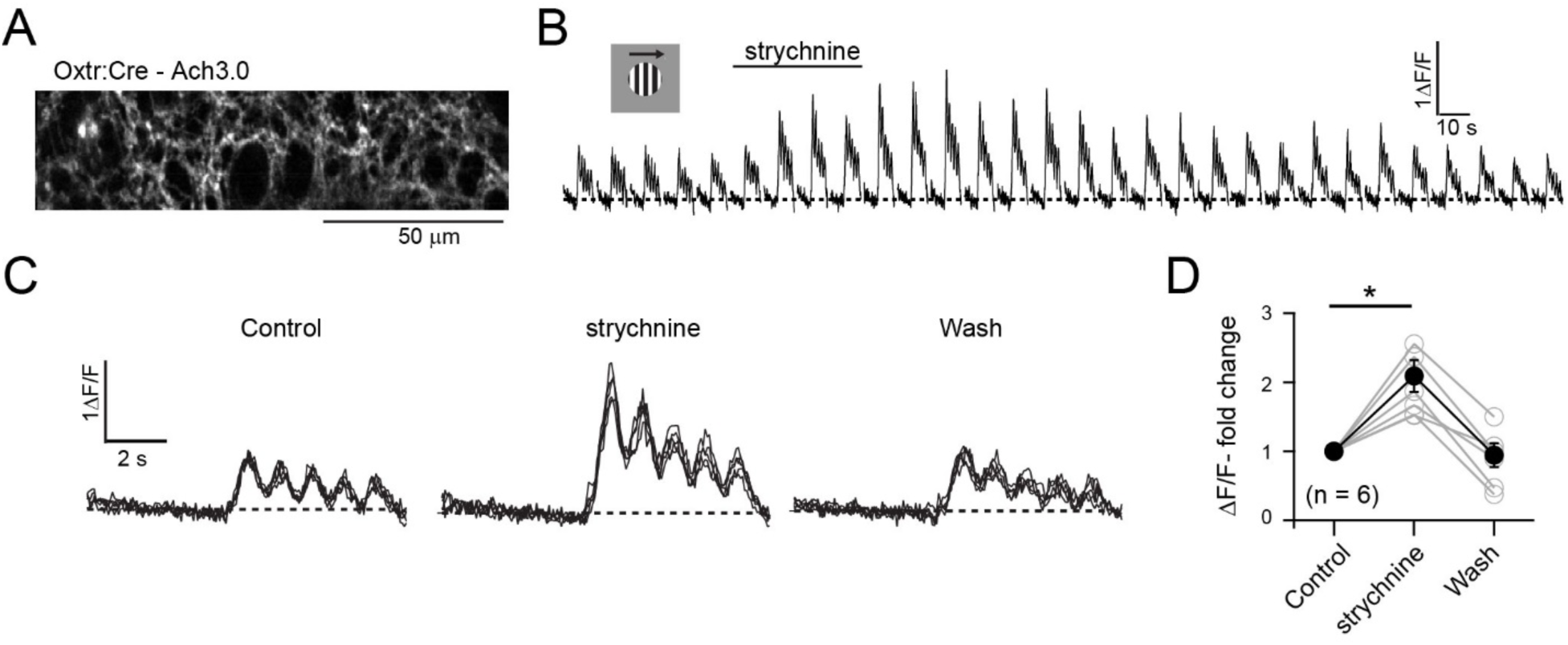
Glycinergic modulation of starburst cholinergic output. **A.** 2P image stack showing the expression of ACh3.0 in dendrites of starburst/DSGC dendrites labelled in the Oxtr-Cre. **B.** The effects of a brief bath application of 1 μM strychnine on the ACh3.0 responses evoked by drifting gratings (SF 0.1cycles/degree; TF 1Hz; duration 5 seconds; presented at 1 minute intervals). The line on top indicates the duration for which the tissue was exposed to strychnine. **C.** Selected ACh3.0 responses shown in **B**, under control (left), strychnine (middle) and recovery from strychnine (wash; right). **D.** Relative change in ACh3.0 fluorescence under the conditions shown in **C**, in 6 field-of-views from 5 retinas (*p = 0.0053). Data represented as mean ± SEM.

### Glycinergic inhibition ensures that SACs maintain direction selectivity throughout their response range

While previous experiments demonstrate that NACs control SAC gain, they do not provide a clear indication of whether or not they affect SAC direction selectivity. To address this question, we examined the effect of strychnine on the direction-selective IPSCs in measured in superior coding Hb9-GFP DSGCs, which receive their inhibitory input primarily from SACs (note, other DSGCs may to receive both SAC and non-SAC input; Park et al., 2015; Sethuramanujam unpublished observations). SAC DS output was quantified by computing the ratio of the peak IPSCs evoked by gratings, drifting in the DSGC’s ‘preferred’ and ‘null’ directions. We found that blocking GlyRs increased inhibitory responses in the null direction and preferred direction, which resulted in weaker direction selectivity. However, this effect was more prominent at non-optimal spatiotemporal frequencies (**Figure 6**).

**Figure 6:**
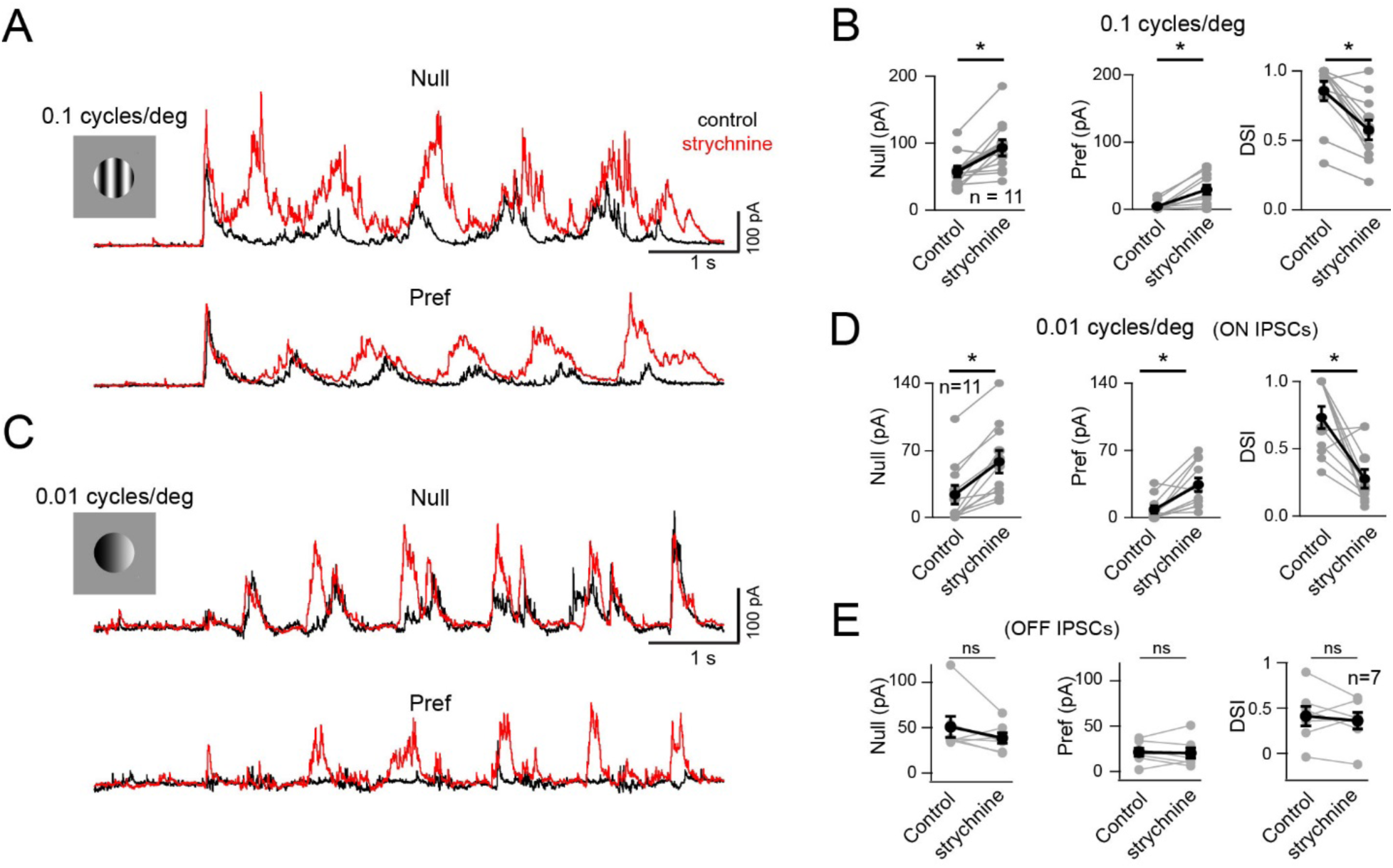
Glycinergic inhibition affects SAC DS output. **A**. IPSCs measured in superior coding DSGCs evoked by gratings (SF 0.1cycles/degree, TF 1Hz) drifting in the null (top) or preferred directions (bottom)(control, black; strychnine, red). For high frequency gratings the ON and OFF components of IPSCs could not be separated. **B.** Average currents evoked by preferred and null stimuli, and the associated direction selectivity index (DSI = (PD-ND)/ (PD+ND), measured across the population (n = 11). **C**. Same as **A**, except gratings were of lower spatial frequency (0.01cycles/degree). Note the weak ON responses in control (black), increase in strychnine for both null and preferred direction stimuli. **D.** Average current and DSI during the ON responses (n = 11). **E.** Same as D, except for OFF IPSCs. Note that on average the OFF IPSCs did not change significantly following strychnine application. Data represented as mean ± SEM.

For example, gratings with low spatial frequency (0.01 cycles/degree) where ON inhibitory currents are weak (**Figure 6C& D**), application of strychnine increased responses in both null (control 25 ± 10 pA and strychnine 59 ± 12 pA; n = 11; p < 0.001) and preferred directions (control 8 ± 4 pA and strychnine 34 ± 7 pA; n = 11; p < 0.005). This resulted in a significant loss of directional inhibition (DSI: control 0.73 ± 0.08 and strychnine 0.28 ± 0.07, p < 0.005; **Figure 6 C&D**). By contrast, the OFF IPSCs in DSGCs remained unaltered in both strength and directionality following glycine receptor blockade. For gratings with high spatial frequency (0.1 cycles/degree), the changes in direction selectivity were less dramatic, as null inhibition was still ∼4X larger than the preferred response (DSI control 0.86 ± 0.07; strychnine 0.58 ± 0.07, p < 0.005) **(Figure 6A&B)**. Although the ON and OFF components of inhibitory currents could not be distinguished at high spatial frequencies, given that strychnine did not affect the OFF pathway, the observed effects are likely to be mediated at the level of ON SACs.

Finally, we directly evaluated the impact of NAC inhibition on SAC direction selectivity (DS), using two-photon Ca^2+^ imaging (**Figure 7A**). We tested the effects of strychnine on the responses of large drifting gratings (0.01 cycles/deg), where their effects are maximum. Consistent with previous reports (Euler et al., 2002; Hausselt et al., 2007), stronger Ca^2+^ responses were observed during centrifugal motion (soma-to-dendrite) compared to centripetal motion (**Figure 7B, C & F**). However, application of strychnine strengthened the centripetal response, significantly decreasing DS Ca^2+^ responses in SAC varicosities (**Figure 7B, C& F**).This effect was more prominent for grating stimuli. When direction selectivity of Ca^2+^ responses was probed with a moving spot, the effect of blocking GlyRs was not significant (**Figure 7D-F**).

**Figure 7:**
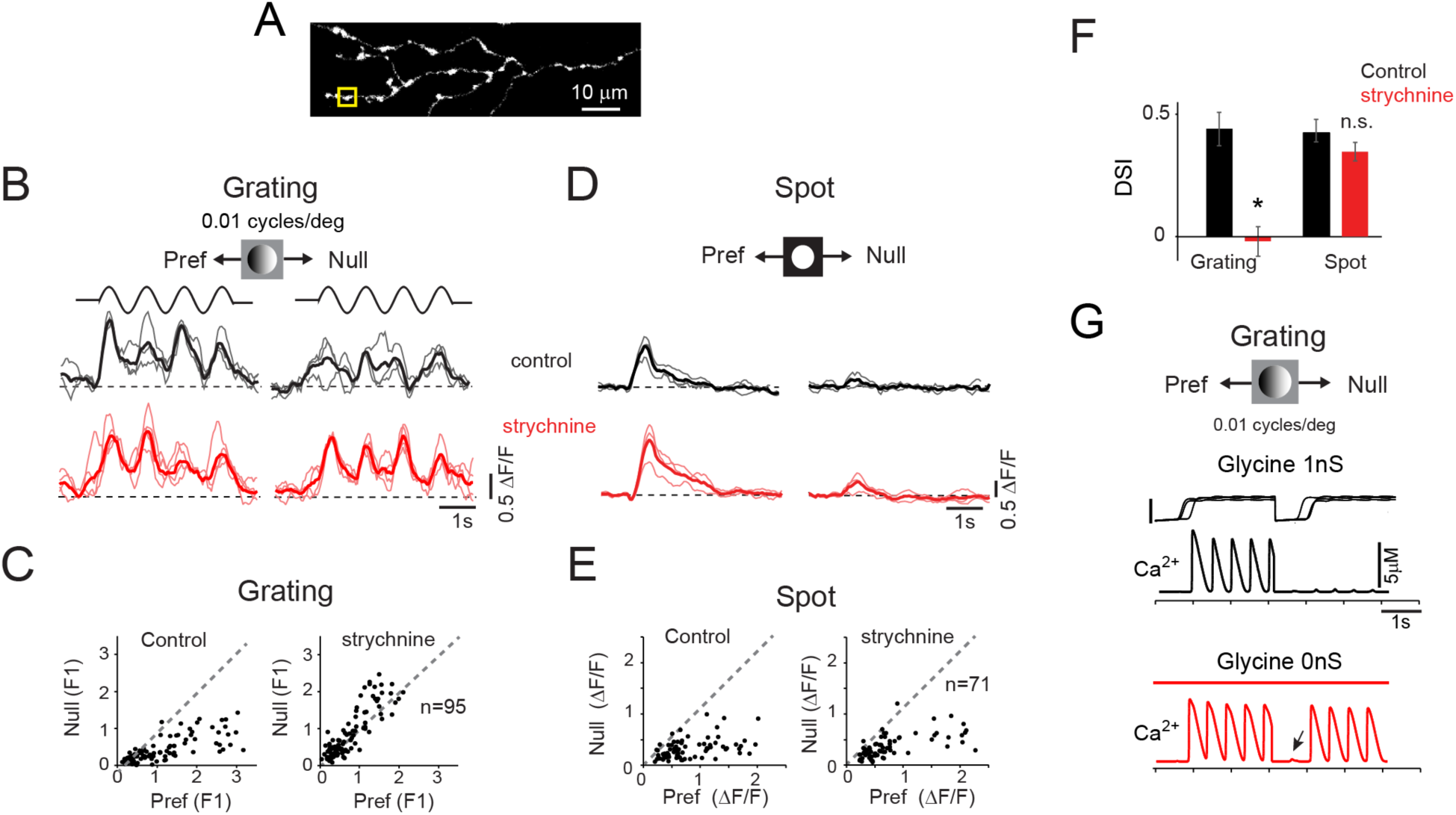
Glycine inhibition is required for the maintenance of DS in SAC dendrites during sustained activity. **A.** Two–photon image stack of ON SAC dendrites filled with Oregon green BAPTA-1 (Ca^2+^ indicator dye). **B.** Change in fluorescence measured at a single distal varicosity (yellow box in **A**) evoked by sinusoidal gratings, moving in the preferred (centrifugal) and null (centripetal) directions, under control (black) and in the presence of strychnine (red). Thicker line in each panel represents the average response of the trials shown. **C**. The *Δ*F/F (F1 component, averaged over 3 trials) measured in individual ROIs for responses evoked in the null direction are plotted against those measured in the preferred direction (Pref), under control (left) and strychnine (right; n = 95 ROIs from 6 ON SACs). **D.** Same as B, except for a 200 μm moving spot. **E.** Change in peak fluorescence for a 200 μm spot moving in preferred and null direction, plotted against each other. Data is pooled from 71 ROIs across 6 cells. **F.** DSI calculated for gratings and spot under control and strychnine. Note the loss of directional Ca^2+^ responses for gratings but not for spots. Data represented as mean ± SEM. **G.** Calcium influx at a distal varicosity of a model ON SAC for a drifting grating in the presence (black) and absence (red) of sustained glycine conductance. Note the directional Ca^2+^ response for the first cycle in absence of glycine inhibition (arrow).

To better understand how glycine affects SAC direction selectivity, we extended the computational models previously used to explore SAC-SAC interactions (Ding et al., 2016) to include the NAC-SAC circuitry. In this model, non-linear processing of inputs by regenerative voltage-dependent Ca^2+^ channels play a key role in amplifying direction selectivity, within a restricted range of SAC EPSP amplitudes (Hausselt et al., 2007; Tukker et al., 2004). Recruiting a sustained glycinergic inhibition from an array of NACs limited the depolarization of the distal SAC dendrites and kept the response within the correct operating range, enabling SACs to produce robust DS (∼0.9) (**Figure 7G**). In the absence of NAC inhibition, however, the null and preferred dendritic responses were no longer in an optimal range, producing similar calcium transients in both centrifugal and centripetal direction (**Figure 7G**), leading to a loss in DS (DSI <0.3). Therefore NAC-mediated gain control appears to be required to gate SAC output, enabling Ca^2+^ channels to be preferentially recruited during strong, but not weak depolarizations.

## Discussion

### Glycinergic NAC inputs: the final pieces of the DS circuit

Since the first evidence that glycine inhibition played a role in the DS circuit (Cunningham and Neal, 1983), several studies have noted specializations in the ON SACs that would suggest that DS may rely on glycinergic inhibition mediated by NACs (Heinze et al., 2007; Ishii and Kaneda, 2014; Majumdar et al., 2009). Here, we directly demonstrate that the glycinergic inputs comprise the major component of light-evoked synaptic inhibition in ON SACs and are critically required to maintain direction selectivity under certain stimulus conditions. This conclusion contrasts with those of a number of previous studies characterizing the anatomical and functional properties of the DS circuitry that suggested a minor role for the NAC pathway.

One reason that NAC function was not considered significant was the fact that they make few contacts with ON SACs (∼5%) and almost no contacts with OFF SACs (Ding et al., 2016). Here we demonstrate several specializations of the NAC pathway that enable them to mediate a powerful inhibition in ON SACs despite their meagre anatomical representation. First, glycinergic transmission at NAC-SAC synapse occurs on an ultra-slow time scale. Second, ON/OFF NACs provide glycinergic inhibition at both phases of a grating stimulus. Third, NAC axonal arbors make well defined mosaics, ensuring that glycinergic inputs could be uniformly activated, throughout the SAC’s receptive field. Together these factors enable signals arising from a few synapses to summate and mediate a sustained inhibition, which is more powerful than that mediated by 95% of the GABAergic synaptic contacts. However, why ON and not OFF SACs require a NAC inhibition for their computations is still not clear.

Another major reason why glycinergic inhibition has been overlooked in the DS circuit is that previous pharmacological studies did not demonstrate significant effects of strychnine on direction selectivity, both at the level of SACs or DSGCs (Caldwell et al., 1978; Wyatt and Day, 1976). However, here we argue that these differences may arise because NAC inhibition is important under specific stimulus conditions. For example, one of the first studies examining SAC DS function showed no effects of strychnine (Hausselt et al., 2007). However, in this study, the stimuli used consisted of expanding/contracting circular rings of a fixed, optimal spatiotemporal frequency that were presented through a “doughnut-shape” window. Since NACs receptive fields are centered around the SAC soma (**Figure 2D**), in retrospect, the minor effects of strychnine on SAC Ca^2+^ responses are not surprising. It is also important to note that NAC inhibition does not modulate responses to simple moving spots (Figure 2E), which are widely used to probe DS mechanisms. Moreover, we found that the effects of blocking NACs on SAC direction selectivity is most robust for sustained stimulation at non-optimal frequencies. Thus, NAC inhibition appears to ensure that SACs maintain DS output throughout their operating range.

At the level of DSGCs, realizing that SAC function is altered upon NAC blockade is complicated because here direction selectivity is shaped by both SAC DS inhibition and the specific SAC-DSGC wiring pattern that create anisotropic GABA/ACh timing offsets (Hanson et al., 2019; Schachter et al., 2010). In addition, DSGCs may also receive glycinergic inputs (Sivyer et al., 2019). Increasing SAC gain upon NAC blockade is not expected to strongly affect direction selectivity in DSGC, because if GABA inhibition and ACh excitation increase in parallel, the computation would remain intact. Moreover, even if SAC output is rendered non-directional, the timing offsets alone may generate robust DS responses in DSGCs. Thus, although blocking GlyRs may have a strong effect on SAC output and render them less DS, owing to the redundancy in DS mechanisms and other network mechanisms the effects of NACs on DSGC responses are expected to be more subtle (**Supplementary Figure 4**).

### Complementary roles for three amacrine cells in the DS circuit

Starburst amacrine cells have the remarkable ability to compute direction of moving objects over a wide range of stimulus spatiotemporal frequencies, likely through engaging multiple DS mechanisms. Reciprocal GABAergic inhibition arising from neighboring SACs plays an important role in shaping the directional tuning properties of SAC dendrites (Chen et al., 2016; Ding et al., 2016; Lee and Zhou, 2006; Munch and Werblin, 2006; Poleg-Polsky et al., 2018) but see (Euler et al., 2002; Hausselt et al., 2007). SAC dendrites are arranged in an ‘anti-parallel’ manner, such that dendrites selective for opposite directions inhibit each other proximal to the soma, where excitatory inputs from bipolar cells are made (Ding et al., 2016; Lee and Zhou, 2006). This ensures that moving objects only activate SAC-mediated surround inhibition on one side of a SAC dendritic field, the side from which they enter, as demonstrated in our electrophysiological recordings (**Figure 2E**). Thus, SAC-mediated inhibition cooperates with other intrinsic DS mechanisms, ensuring that objects moving from the dendritic tip to the soma (centripetal motion) produce weak responses. Reciprocal GABAergic inhibition is especially important when stimulus contrast is high, where it ensures that responses to strong excitatory inputs do not drive depolarizations during non-preferred centripetal motion (Ding et al., 2016). It is important to note that SAC-mediated GABAergic inhibition only occurs when SACs themselves are strongly stimulated.

However, NACs are driven by a semi-independent pathway mediated by an overlapping set of ON/OFF bipolar cells. This could enable NACs to provide inhibition over a significantly broader range of contrast, compared to SACs. Moreover, SAC output was enhanced in both preferred and null directions by glycinergic block, indicating that the glycinergic inhibition itself is non-directional. Importantly, direction selectivity in SAC dendrites was significantly degraded when NAC inhibition was blocked, especially for low spatial frequency gratings. Under these conditions SAC-SAC GABAergic inhibition is augmented, but yet not sufficient to prevent the SAC from becoming less DS. Thus, the powerful NAC-mediated glycine inhibition appears to be well-suited and critically required for optimal DS SAC output under continuous stimulation patterns.

Like NACs, WACs provide inhibition that is also non-directional and shapes the tuning properties of SACs, but in a distinct way. For example, WACs control the spatiotemporal selectivity, without altering the DS response of the SAC (Hoggarth et al., 2015; Huang et al., 2019). In contrast, NACs do not integrate information over large areas, but strongly impacted DS processing in SACs, as discussed above. Interestingly, NACs and WACs are likely to be reciprocally connected (Berson, unpublished results; also see (Werblin, 2010)). Our results do not address this question, as our focus was to understand the properties of NAC-SAC connections and stimuli were presented through an aperture that minimize the surround inhibition mediated by WACs. Future investigations are required to probe NAC-WAC interactions to directly determine how these three distinct amacrine cell types interact with each other to enable direction coding under natural stimulus conditions.

### Glycinergic inhibition shapes direction selectivity in SAC dendrites

The mechanisms by which glycinergic inhibition impacts SAC function can be best understood in the context of existing models proposed for SAC DS. These suggest that SACs rely on differences in temporal summation of EPSPs to produce directional responses. Specifically, from the perspective of their distal dendrites (where output occurs), optimal summation occurs for centrifugal motion. Importantly, these anisotropies in temporal summation are strongly amplified by voltage-gated Ca^2+^channels, in a manner that is steeply dependent on voltage. Thus, in these models an effective gain control mechanism is a key requirement, to keep the dendritic response within a restricted range, regardless of the strength of the stimulus. Without gain control, the mechanisms underlying DS would easily saturate. Reciprocal inhibition within the SAC network has been previously proposed to serve as a source of gain control (Ding et al., 2016). Consistent with this hypothesis, blocking GABA receptors significantly affects ability of SACs to compute DS when stimulated with high-but not low-contrast spot stimuli (Ding et al., 2016). However, in the context of sustained activity, like that driven by drifting grating stimuli, we find applying strychnine dramatically increases the SAC GABA output and compromises the ability of starburst dendrites to compute direction. Thus, even the resulting increased level of gain control offered by the SAC network does not appear to be adequate for maintaining DS under sustained stimulation. This is likely because during continuous stimulation there is a significant ‘DC’ component, which cannot be easily cancelled by a transient form of SAC inhibition. For canceling the sustained excitatory components, slow-glycinergic inhibition mediated by NACs appears better suited. By strongly shunting SAC dendrites and lowering their input resistance, the NAC inhibition to SACs lowers the gain of SAC dendritic EPSPs and, thus allows SAC dendrites to more effectively compute directional differences without the saturating effects of DC signals. In addition, the strong glycinergic shunt near the soma may also help preserve the electrical isolation of individual SAC dendrites. This would prevent excitation from spreading throughout the arbor, which would degrade its ability to compute direction. However, the precise degree to which this glycinergic input helps in electrical isolation remains to be assessed experimentally.

### Other synapses expressing Glyα4Rs subtype

Slow inhibition appears to be a function of specialized GlyRs containing α4 subunits, as spontaneous and light-evoked slow IPSCs were mostly absent from SACs in the *Glra4* KO mouse. However, in a few SACs we did note residual GlyR-mediated spontaneous IPSCs suggesting that other subunits such as GlyRα2 may also participate in SAC function (Heinze et al., 2007; Majumdar et al., 2009). ON DS ganglion cells have been recently reported to receive both fast and slow forms of postsynaptic glycinergic inhibition, but which GlyR subtypes and NACs are involved remains unknown (Sivyer et al., 2019). Nevertheless, synaptic transmission at other α4-containing glycinergic synapses is also consistent with our conclusions. Some of the WACs and other types of NACs (non-AII ACs) are reported to express α4 receptors, and exhibit slow kinetic spontaneous events (decay constant τ ∼30ms) in conjunction with α2 glycine receptors (Heinze et al., 2007; Majumdar et al., 2009). Knocking out GlyRα2 receptors increased the *τ*_decay_ of IPSCs to ∼70 ms (similar to what we have measured in ON SACs), providing further evidence for the role of α4 in slow synaptic inhibition (Weiss et al., 2008). Furthermore, expression studies demonstrate that mouse α4β receptors mediate slow sIPSCs (*τ*_decay_ ∼80ms) (Leacock et al., 2018), comparable to the decay of sIPSCs we measured in SACs. Slow transmission due to paracrine/extra-synaptic mechanisms is also a possibility. However, the majority of GlyRs in retina express β subunits, which localizes them at synapses through interactions with gephyrin (Fischer et al., 2000; Meyer et al., 1995). Thus, it is likely that the slow properties of NAC-SAC synapses arise from the biophysical properties of GlyR α4.

### Conclusions and future directions

In summary, our results demonstrate that NACs likely play an important role in mediating DS responses in SACs and do so in a way that is distinct from other forms of inhibition mediated by SACs and WACs, highlighting the role of multiple interneurons within a given neural circuit. It is interesting to note that although the GlyRs mediating NAC responses are ultra-slow, they participate in a rapid computation. Interestingly, in zebrafish, GlyRα4 appears to be involved in rapid escape and startle responses (Leacock et al., 2018), raising the possibility that tonic inhibition may be used to maintain computational efficacy in other circuits. Thus, neural circuits appear to employ interneurons with diverse anatomical and functional properties for their complex computations.

## Acknowledgements

We thank Tracy Michaels for performing AAV injections and help with mouse colony management. Dr. Yulong Li & Dr. Miao Jing (Peking University) for providing genetically encoded ACh sensor. Dr. Rudolph Uwe (Harvard Medical) for Gabra2 KO mice; Dr. Marla Feller for nGFP mice. Dr. Jamie Boyd for his help with IGOR software for 2P imaging. This work was supported by CIHR (GBA); NIH (RGS, EY022070; MAMc, EY14701 and 29719; DB EY14701); and Kentucky Lions Eye Research Endowed Chair (MAMc).

## Author Contributions

VJ: Data acquisition, data analysis, writing—original draft, review, and editing. LH: Ca^2+^ imaging, spike recordings, data analysis, review and editing. SS: ACh imaging, spike recordings, data analysis, review and editing. RGS: Computational modeling, review and editing. MAMc generated and tested Glra4 KO mouse, review and editing; RGG, CZ generated and tested Glra4 KO mouse, DB: SBEM analysis, review and editing. GA: supervision, funding acquisition, experimental design, writing—review and editing.

The authors declare no competing interests.

## Supplementary Figures

**Supplementary Figure 1.**
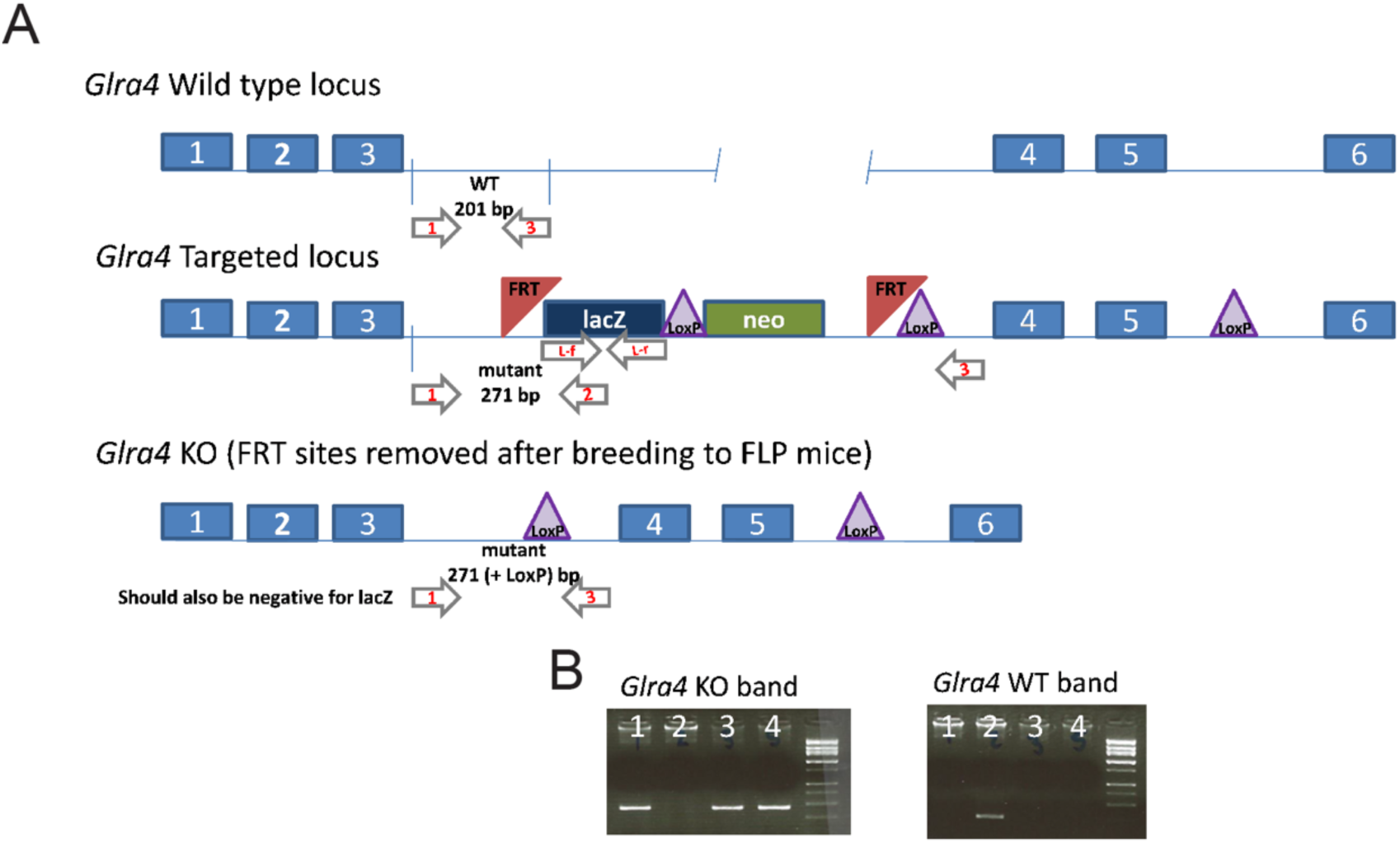
Targeted deletion of GlyRα4 and generation of *Glra4* KO mice. **A.** The wild-type mouse locus of Glra4, its targeted locus and the production of a *Glra4* KO after breeding to FLP mice to remove all DNA, between and including the FRT sites in the targeted locus. **B.** PCR products from primers 1 and 3 show that animals 1, 3 and 4 are KO mice and animal 2 is a wild-type littermate. NOTE *Glra4* is located on the X chromosome, which makes it possible to obtain Glra4 KO males and wild-type females in the same litter.

**Supplementary Figure 2.**
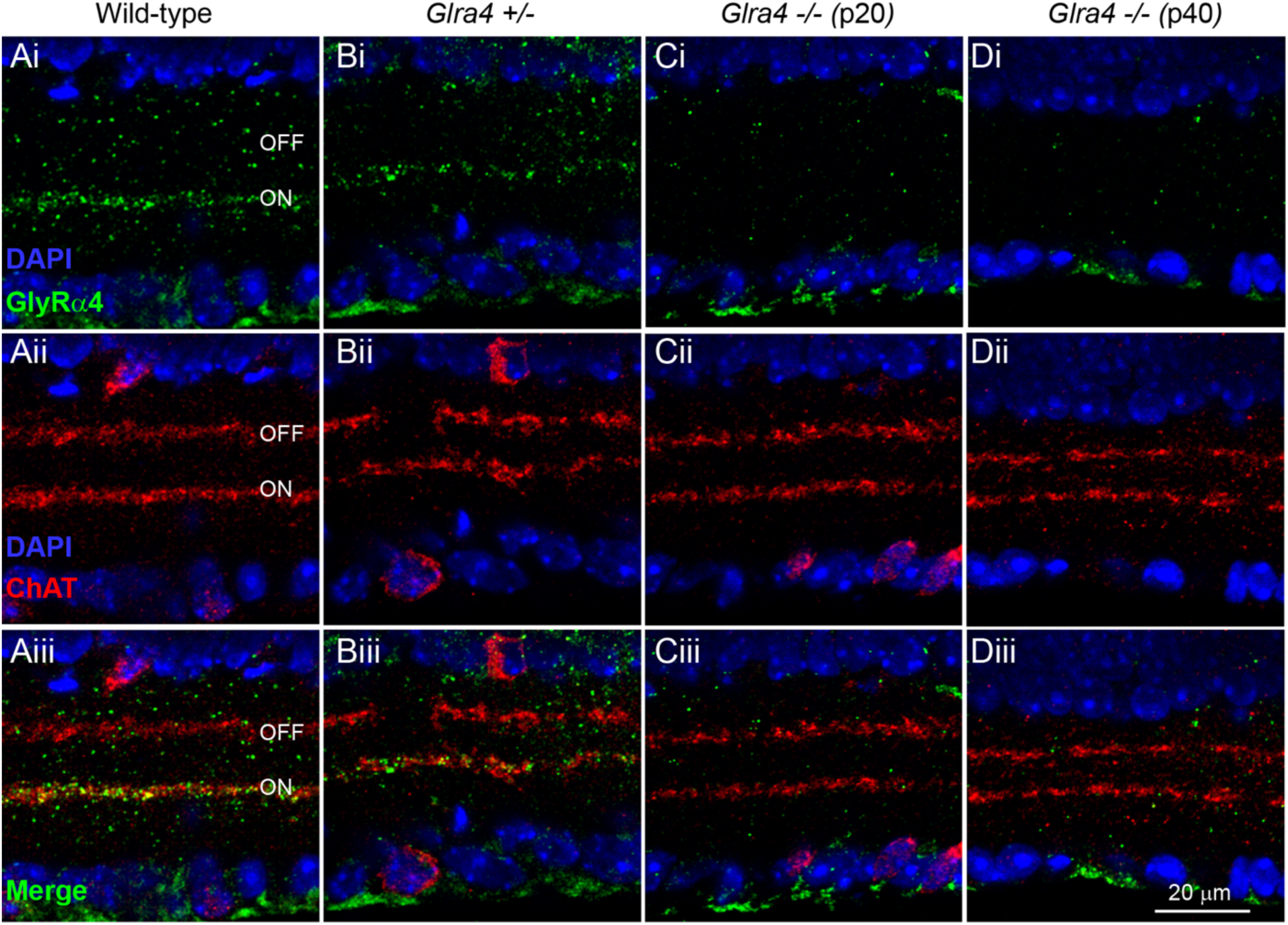
GlyRα4 expression is absent in *Glra4* KO mice. **A.** Confocal images of a wild-type mouse retinal section reacted with anti-GlyRα4 (i) and anti-cholineacetyl transferase (ChAT, ii). The merged image (Aiii) clearly illustrates the co-localization of the expression of the two proteins, primarily in the ON ChAT band of the inner plexiform layer (ON). **B.** A similar expression profile is seen in a mouse heterozygous for *Glra4*^+/-^. **C & D**. In both young (Postnatal day (P) 20 and mature (P40) retinas of *Glra4* -/- mice, expression of GlyRα4 is absent and has no effect on the expression of ChAT. Scale bar for all images 20 µm.

**Supplementary Figure 3.**
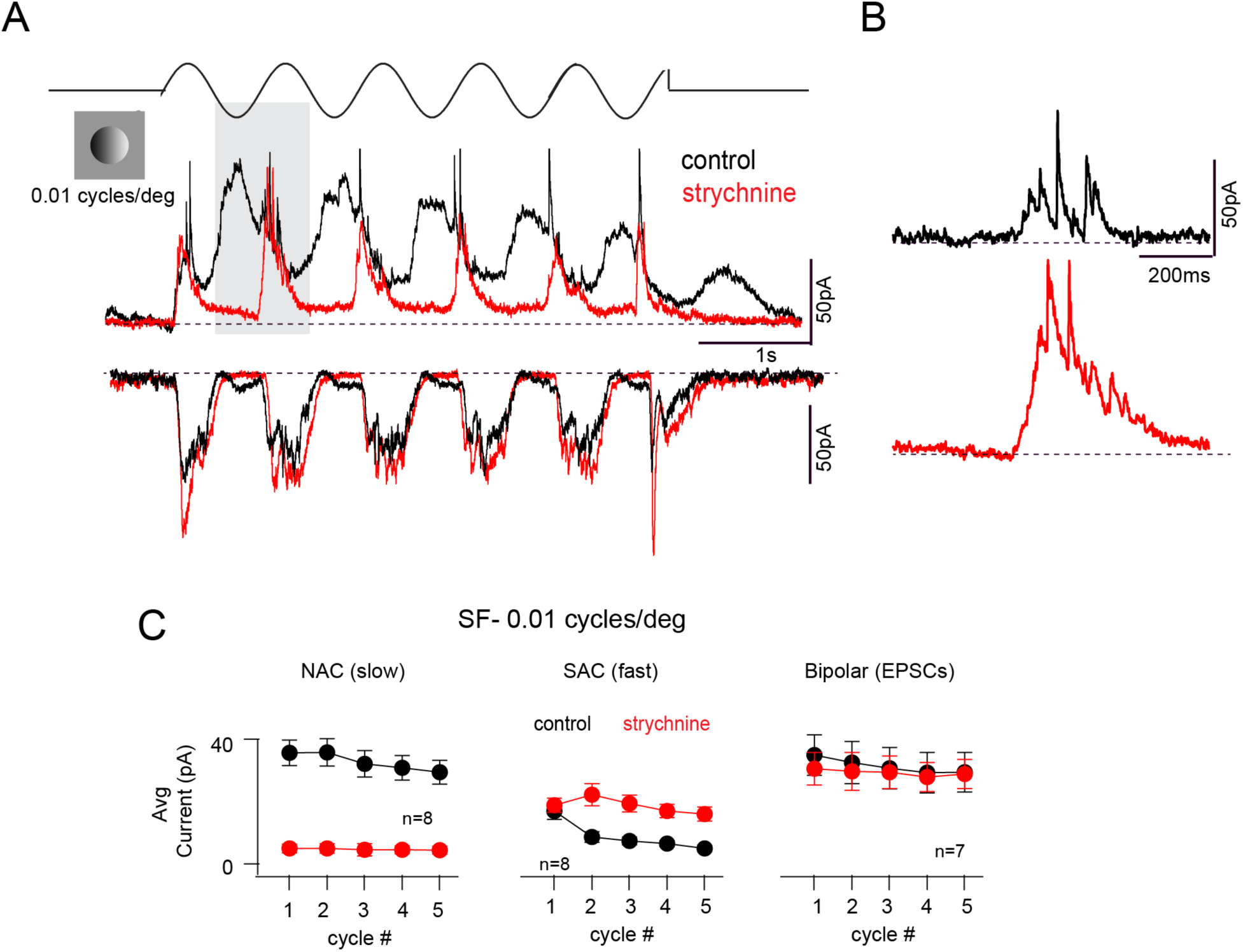
Suppression of SAC component at low spatial frequency. **A.** IPSCs (0mV) and EPSCs (−60mV) measure from ON SACs under control (black) or in the added presence of strychnine (red). Grating SF 0.01cycles/degree, TF 1Hz. **B.** Shaded region in A magnified to show the fast SAC-mediated transient IPSCs are enhanced in the presence of strychnine. **C.** Strychnine has distinct effects on the NAC-, SAC-mediated IPSCs (middle) and EPSCs at each of the 5 cycles of the stimulus (control, black; strychnine, red). Data represented as mean ± SEM.

**Supplementary Figure 4.**
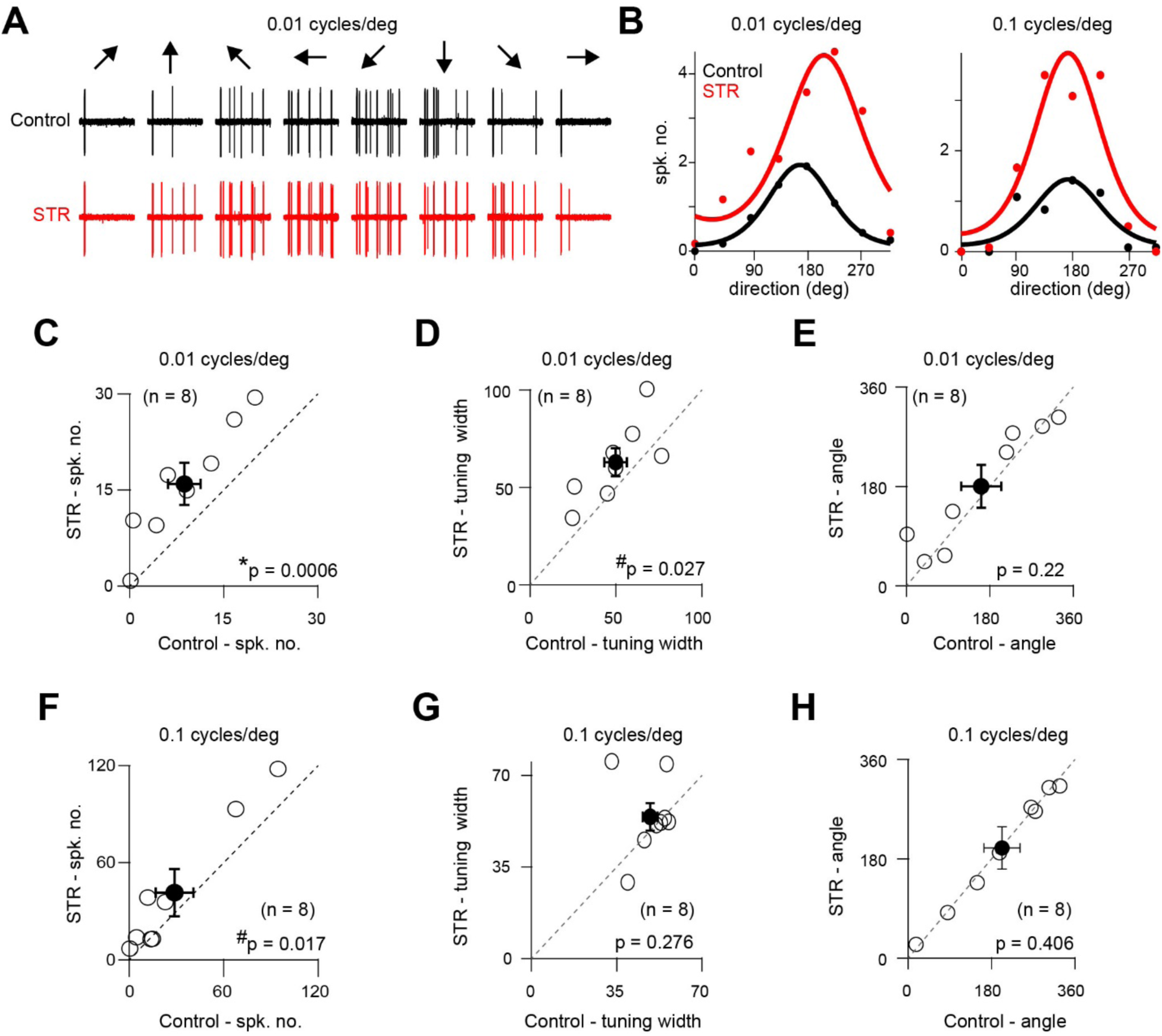
GlyR block increases DSGC spiking responses but does not abolish direction selectivity. **A.** Spike activity in DSGCs elicited to gratings drifting in 8 different directions (arrows on top), under control (black) and in strychnine (red). The data illustrated here is for gratings with SF 0.01cycles/deg (TF 1Hz). **B.** Tuning curves for directional spiking responses generated in a DSGC in response to drifting gratings with SF 0.01cycles/deg (left) and 0.1 cycles/deg gratings (right). Responses in both control and strychnine are shown. These responses were measured as the average number of spikes per cycle (first cycle was excluded from the analysis). The line shows the data fitted to a Von Mises function (See methods). **C – E.** The amplitudes (C), tuning widths (D) and preferred directions (E) of DSGC responses generated by drifting gratings of SF 0.01cycles/deg, in control and strychnine (n = 8; paired t-tests). The tuning width and preferred directions were estimated from the Von Mises fit (See methods). **F – H.** Similar to C – E, but for drifting gratings of SF 0.1cycles/deg. Data represented as mean ± SEM.

## Methods

### Animals

Experiments were performed using adult (P30 or older) mice of either sex: C57BL/6J (JAX:000664). In most experiments, SACs and DSGCs were identified in mouse lines in which they were fluorescently labelled, using 2-photon imaging. Superior coding DSGCs were targeted in Hb9^eGFP^ (RRID: MGI_109160) line, in which they are fluorescently labelled. ON SACs were genetically accessed using the choline acetyltransferase (ChaT) cre-mouse line (RRID: MGI_5475195). To label SACs with fluorescent markers the ChaT-cre line was crossed with Ai9 (RRID:MGI_3809523); or floxed nGFP line (Kindly provided by Dr. Marla Feller, University of California at Berkeley). To reduce SAC GABA release the ChaT-cre line was crossed with a floxed allele of Slc32a1 (*Slc32a1^tm1Lowl^*, JAX stock # 012897), which also is commonly referred to as *Vgat^flox/flox^;* JAX012897). To block SAC-SAC GABA transmission postsynaptically, the ChaT-cre line was crossed with *Gabra2^tm2.2Uru^*. Glrα4 KO mice were generated from ES cells purchased from the KOMP Repository (*Glra4^tm1a(KOMP)Wild-types^*; RRID:MMRRC_056080-UCD), and used by the Transgenic Core Facility at Cincinnati Children’s Hospital Medical Center to generate chimeric mice. Supplemental Figure 1 shows the WILD-TYPE and targeting locus for *Glra4*. Founder mice were crossed to C57BL6/J mice to produce heterozygous animals carrying the targeted allele. To generate the *Glra4* KO used in this study mice carrying the KOMP allele were crossed to *B6N.Cg-Edil3^Tg(Sox-cre)1Amc^* (Jax stock # 014094), and offspring carrying the floxed allele were identified by PCR. This removed all of the targeting construct between the LoxP sites, including exon 3. The genotype of the progeny were determined using PCR and the following primers, which differentiated the wild-type allele (201 bp) from the KO allele (271 bp).

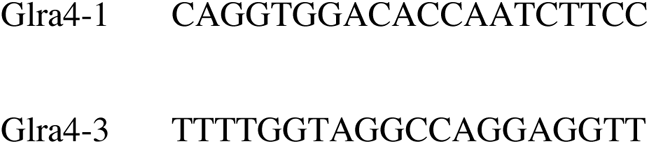

Because *Glra4* is located on the X chromosome, female heterozygous mice were bred to C57BL/6J males and the first generation produced *Glra4* homozygous males. The progeny were backcrossed onto C57BL/6J for 8 generations and the animals then maintained as homozygous in the breeding colony thereafter. To characterize the changes in GlyRα4 expression, we use a previously published immunohistochemistry protocol (Nobles et al., 2012) on 20 µm cryostat sections. We examined and compared the expression pattern of GlyRα4 in wild-type, heterozygous and KO mouse retinal section (**Supplemental Figure 2**), using a rabbit anti-Glra4 antibody (1:100; Chemicon Cat# AB9696) with a goat anti-rabbit Cy3 secondary (1:200, Jackson ImmunoResearch Cat# 705165003). We examined the pattern of GlyRα4 relative to the bands formed by the ON and OFF SAC dendrites in the IPL, using a goat anti-Choline Acetyltransferase antibody (1:200; Chemicon Cat# AB144) with a donkey anti-goat IgG Alexa 405 (1:200, Abcam Cat# ab175664). Confocal images were acquired on an Olympus FV1000 confocal microscope (Olympus, U.S.A.) and z-stack images were acquired with 0.2 μm z-steps using a 63x oil immersion lens (NA 1.4). In wild-type and heterozygous mouse retina sections, GlyRα4 expression is found only in the IPL and is most highly concentrated within the band formed by the ON SAC dendrites (immunopositive for ChAT) (**Supplemental Figure 2**). The developing (P20) and mature (≥ P40) *Glra4* KO retinal sections do not express GlyRα4. We attempted to use this same antibody in Western blots and found that a band of the same size is labeled in wild-type and Glra4KO retinal fractions, thus this antibody is not useful for detecting GlyRα4 in Western blots (data not shown). All procedures were performed in accordance with the Canadian Council on Animal Care and approved by the University of Victoria’s Animal Care Committee or the University of Louisville Animal Care and Use Committee.

### Retinal preparation and physiological recordings

Mice were dark-adapted for at least 1 hour before being anesthetized with isoflurane and decapitated. Retinas of both eyes were extracted in standard Ringer’s solution under a dissecting microscope equipped with infrared optics. Isolated retinas were laid flat, ganglion cell side up, over a pre-cut window in 0.22 mm membrane filter paper (Millipore). Retinas were then placed in a recording chamber maintained at 35°C and continuously perfused with Ringer’s solution (110 mM NaCl, 2.5 mM KCl, 1mM CaCl_2_, 1.6 mM MgCl_2_, 10 mM glucose, 22 mM NaHCO_3_) that was bubbled with 95% CO_2_/5% O_2_. Retinas were visualized using a BX-51 WI microscope (40x water-immersion objective; Olympus, Canada) with the help of an infrared CCD camera (Spot RT3 Diagnostic Instruments). Patch-clamp recordings were made with a MultiClamp 700B amplifier (Molecular Devices). Patch electrodes (5-8 MΩ) contained the following (in mM): 112.5 CH_3_CsO_3_S, 7.75 CsCl, 1 MgSO_4_, 10 EGTA, 10 HEPES, 5 QX-314-bromide (Tocris). The pH was adjusted to 7.4 with CsOH. The reversal potential for chloride was calculated to be - 56mV. The recordings were not corrected for series resistance or the junction potential. Signals were digitized at 10 kHz (PCI-6036E acquisition board, National Instruments) and acquired using custom software written in LabVIEW. Data was analyzed offline with custom written software in MATLAB. Unless otherwise noted, all reagents were purchased from Sigma-Aldrich Canada. TTX and gabazine were purchased from Alamone Labs and Tocris, respectively.

### Virus injections

The single-stranded AAV vector ssAAV-9/2-hSyn1-chI-dlox-Igk_(rev)-dlox-WPRE-SV40p(A) (1.2 × 10^13^ vg/ml) was produced using the plasmid by Viral Vector Facility at University of Zurich (Addgene #121922). For intravitreal viral injections of the AAV, mice were anesthetized with an i.p. injection of fentanyl (0.05 mg/kg body weight; Actavi), midazolam (5.0 mg/kg body weight; Dormicum, Roche) and medetomidine (0.5 mg/kg body weight; Domitor, Orion) mixture dissolved in saline. We made a small hole at the border between the sclera and the cornea with a 30-gauge needle. Next, we loaded the AAV into a pulled borosilicate glass micropipette (30 *µ*m tip diameter), and 2 *µ*l was pressure-injected through the hole into the vitreous of the left eye using a Picospritzer III (Parker). Mice were returned to their home cage after anesthesia was antagonized by an i.p. injection of a flumazenil (0.5 mg/kg body weight; Anexate, Roche) and atipamezole (2.5 mg/kg body weight; Antisedan, Orion Pharma) mixture dissolved in saline and, after recovery, were placed on a heating pad for one hour.

### Two-photon functional imaging

For ACh3.0 sensor imaging, experiments were performed three-to-four weeks after virus injection into the eyes of Oxtr-T2A-Cre or CART-Cre mice (to label SAC/DSGC dendrites). For calcium imaging, SAC somas labelled in ChaT-cre x nGFP mouse line were electroporated with a calcium indicator Oregon Green 488 Bapta-1 (15mM in water; ThermoFisher Scientific)(Ding et al., 2016). Retinas were prepared following a procedure that was similar to that used for the electrophysiological recordings. SAC/DSGC dendrites were imaged using a custom two photon microscope equipped with Ti/Sapphire laser (Insight DeepSee^+^:Spectra Physics) tuned to 920 nm. The laser was guided by XY galvanometer mirrors (Cambridge Technology) controlled with software developed by Dr. Jamie Boyd (University of British Columbia) in the Igor Pro environment. Image scans were acquired at 10-15 Hz with a XY pixel size of 0.28µm X 0.28µm, and a dwell time of 1µs/pixel. Imaging windows were typically 125µm X 50µm in area.

### Visual Stimulation

Visual stimuli were produced using a digital light projector and were focused onto the photoreceptor layer of the retina through the sub-stage condenser. The background luminance was measured to be 10 photoisomerisations/rod/second (R*/rod/s). Unless mentioned, all the experiments were performed in dark adapted state. Stimuli were designed and generated using Psychtoolbox (MATLAB, Mathworks). Spots were presented on a dark background, while gratings were presented on a grey background (400 R*/rod/s) through a 500 μm circular window. Sine wave gratings with spatial frequency 0.1 and 0.01 cycles/deg moving at 1Hz were used,and were presented once every 10 seconds. Stimulus widths were converted to cycles/degree of visual angle assuming 1 degree ∼30 μm on the mouse retina (Remtulla and Hallett, 1985).

### Data Analysis

#### Electrophysiology

Physiological data were analyzed using custom routines written in Matlab or Igor (Wavemetrics). For stationary spots, ON IPSCs measured in SACs were separated into transient and sustained components based on their timing. The strength of the transient component was estimated integrating IPSC over the first 100 ms of the response, whereas the sustained component was estimated by integrating the rest of the response. For the OFF IPSC, the entire response was considered to be sustained. For moving spots, the temporal windows over which responses were integrated were based on the pharmacological profile of the transient and sustained inhibitory components.

For grating stimuli, for each cycle, first the total charge (area under the curve) was measured. Then, sustained currents were baselined to estimate the transient component, as shown in Fig 4B. Finally, the strength of sustained component was computed by subtracting the transient component from the total charge. For the sustained IPSCs, ON and OFF components could not be easily separated, thus the whole area under the sustained region was considered as the slow current mediated by glycine receptors. In ON-OFF DSGCs, ON and OFF components could not be separated for high spatial frequency gratings (SF 0.1 cycles/degree) but could be separated for lower spatial frequency gratings (SF 0.01 cycles/degree). A photodiode was used to measure the phase of the stimulus.

To estimate the tuning width and preferred direction of the DSGC responses, the data was fitted to a Von Mises fit defined by the following equation;

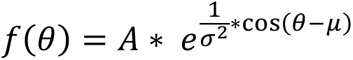

where *θ*indicates the direction of motion, A refers to the amplitude, μ indicates the mean and *σ* the standard deviation. *σ* is indicated as tuning width and μ as the preferred direction in **Supplementary Figure 4**.

Spontaneous IPSCs measured in SACs (V_HOLD_ ∼ 0mV) in dark-adapted retinas and analyzed using Taro tools software in Igor. Only events with amplitudes above 5 pA were selected for analysis. The rise of the sIPSC was estimated by measuring the time taken for the event to rise from 20 to 80% of its maximal amplitude. The decay kinetics were estimated using single exponential function (only non-overlapping events were chosen to measure decay kinetics).

#### Two Photon Imaging

For the ACh3.0 signals, light-evoked changes in fluorescence were measured over the whole field-of-view. In the case of Ca^2+^ signals, changes in fluorescence were measured in individual SAC varicosities. Light-evoked changes is fluorescence (ΔF) were normalized to the baseline fluorescence (F) measured before stimulus onset, corrected for background fluorescence (Fb). In the case of AC3.0 signals, the average ΔF/F over the duration of the grating stimuli was used to quantify the strength of the response. In the case of Ca^2+^ signals, for moving spots, the peak ΔF/F was used to quantify the strength of the response, while for sine wave gratings, the magnitude of the fundamental harmonic was used (F1 component). DSI was calculated as (PD-ND)/ (PD+ND), where PD and ND refer to responses evoked by stimuli moving in the preferred and null direction, respectively.

### Anatomical Reconstruction

A previously published dataset acquired using scanning SBEM was analyzed (retina k0563; (Ding et al., 2016)). Voxel dimensions were 12 × 12 × 25 nanometer (nm) (x, y, and z, respectively). ON-starbursts were identified and back traced to identify the pre-synaptic NACs. Bipolar ribbon synapses were identified on the NACs, by using their synaptic ribbons. All analyses were performed by tracing skeletons and annotating synapses using the Knossos software package (https://knossostool.org/).

### Computational Model

We used a computational model to estimate membrane potential, intracellular Ca^2+^ or local synaptic conductances in SAC dendrites. The model comprised a 7-cell SAC network (1 cell in the center with 6 neighbors in a hexagonal array) incorporating mouse connectivity, utilizing several example morphologies with random rotations to minimize results peculiar to one morphology or orientation (Ding et al., 2016; Stincic et al., 2016). SAC dendritic tips made reciprocal GABAergic connections on their neighbors’ opposing dendrites proximal to the soma. The SAC morphologies were discretized with a compartment size of 0.1 lambda (space constant), giving 170-200 compartments per cell. Bipolar cells were placed by an algorithm in a semi-regular array according to the known regularity of their spacing and based on their proximity made synaptic contact with AMPA receptors on SAC dendrites within a radius of ∼100 *µ*m of the soma(Ding et al., 2016). Biophysical properties (Ri, Rm, channel conductances and densities, synaptic properties and distributions) were set from previous models (Ding et al., 2016; Stincic et al., 2016). During simulated motion, roughly 25 GABAergic SAC-SAC synapses were activated, each with a unitary synaptic conductance that ranged between 25-100 nS, producing a total inhibitory conductance that ranged between 500-1000 nS at the central model SAC. Glycinergic input to SACs came from narrow-field amacrine cells that made a total of ∼20 synaptic connections to each SAC within a 40 *µ*m radius of the soma. Each glycinergic synapse had a maximum conductance of 40-50 nS, contributing a total inhibitory conductance to each SAC of 500-1000 nS. The glycinergic synapses were given a time constant of 500 ms. Kv3 channels were included at the SAC soma (3mS/cm^2^) and proximal dendrites (2mS/cm^2^) for current clamp recordings from the soma but were removed for somatic voltage clamp recordings. Slowly inactivating calcium channels were included in medial and distal SAC dendrites (9 mS/cm^2^) and generated regenerative peaks to −20 mV when SAC dendritic EPSPs were depolarized above −45 mV. Ligand and voltage-gated ion channels were modeled using Markov sequential-state diagrams. Visual signals were simulated in the model’s bipolar cells by voltage clamping them with a spatio-temporal stimulus pattern specified as a resting potential that produced a low continuous vesicle release rate and contrast defined as a depolarization in the range of 1-10 mV. The 7-cell network was stimulated with a sine-wave grating (300 *µ*m/cyc, 600 *µ*m/s) at a background sufficient to depolarize (with the given synaptic input) the SACs to a somatic V_rest_ of −55 to −50 mV with voltage peaks to −40 mV. Models were run with the Neuron-C simulation language on an array of 12 3200 MHz Opteron servers (each with 2 CPUs, 8 cores) running Linux-64 under the Mosix parallel job-sharing system.

